# Early alterations of motor learning and corticostriatal network activity in a Huntington’s disease mouse model

**DOI:** 10.1101/2024.10.07.616936

**Authors:** N Badreddine, F Appaix, G Becq, S Achard, F Saudou, E Fino

## Abstract

Huntington’s disease (HD) is a neurodegenerative disorder that presents motor, cognitive and psychiatric symptoms as it progresses. Prior to motor symptoms onset, alterations and dysfunctions in the corticostriatal projections have been described along with cognitive deficits, but the sequence of early defects of brain circuits is largely unknown. There is thus a crucial need to identify early alterations that precede symptoms and that could be used as potential early disease markers. Using an HD knock-In mouse model (Hdh^CAG140/+^) that recapitulates the human genetic alterations and that show a late and progressive appearance of anatomical and behavior deficits, we identified early alterations in the motor learning abilities of young mice, long before any motor coordination defects. In parallel, e*x vivo* two-photon calcium recordings revealed that young HD mice have altered basal activity patterns in both dorsomedial and dorsolateral parts of the striatum. In addition, while wild-type mice display specific reorganization of the activity upon motor training, network alterations present in the basal state of non-trained mice are not affected by motor training of HD mice. Our results thus identify early behavioral deficits and network alterations that could serve as early markers of the disease.

## INTRODUCTION

Huntington’s disease (HD) is a neurodegenerative disorder characterized by motor, cognitive and psychiatric symptoms, stemming from an expansion of the CAG repeat in exon 1 of the huntingtin gene (Huntington’s Disease Collaborative Research Group, 1993). HD primarily affects the striatum (Vonsattel et al., 1985) followed by certain cortical areas. Even though motor symptoms appear later, cognitive and learning deficits have been widely described ten years prior (Biglan et al., 2016; Papoutsi et al., 2014). Interestingly, the assessment of the speed of processing, initiation and attention measures seem to allow for a better diagnosis of HD and prediction of functional decline (Paulsen, 2011). Indeed, the PREDICT-HD study, using a speeded tapping test, showed a decline of movement speed and motor skills in prodromal HD patients (Stout et al., 2011), suggesting the interest of using motor skill learning in early HD detection.

Pathophysiological studies indicate that disturbances in corticostriatal connections precede late-stage degeneration. Imaging and electrophysiological animal studies reported elevated cortical activity in premanifest HD (Arnoux et al., 2018; Burgold et al., 2019; Donzis et al., 2020). We and others have reported an altered synchrony between cortical and striatal networks (Lee Hong & Rebec, 2012; Naze et al., 2018; Virlogeux et al., 2018), thus suggesting the occurrence of network alterations before the start of degeneration. In an attempt to link dysfunctions in the cortex to the behavioral symptoms observed in premanifest HD, Deng et al. showed that significant reduction of the number of corticostriatal terminals only occurs in (> 12 month-old) knock-in HD mice (Hdh^CAG140/+^), linking thus cortical alterations and motor impairments (Deng et al., 2013). More recently, alterations in the corticostriatal networks early during the first postnatal week, were linked to the behavioral deficits that occur in adulthood (Braz et al., 2022). It is therefore important to identify neuronal markers of early dysfunctions, before the first appearance of marked motor symptoms and eventually how it relates to functionally distinct corticostriatal territories.

Corticostriatal connections are involved in wider cortico-basal ganglia-thalamocortical loops (Redgrave et al., 2010). Within the striatum, the dorsomedial part (DMS) is integrated in the associative loop, and the dorsolateral part (DLS) in included in the sensorimotor loop. In physiological conditions, DMS and DLS have been shown to play distinct roles in striatal-dependent tasks, with a preferential involvement of the DMS in flexible behaviors and action-outcome association, and of the DLS in sensorimotor association and habitual behavior (Balleine et al., 2007; Kupferschmidt et al., 2017; Redgrave et al., 2010; Smith & Graybiel, 2013, 2016; Thorn et al., 2010; D. Yin et al., 2009). Those distinct striatal territories are likely to be differently associated to specific symptoms in HD. Notably, DMS and DLS are not affected the same way in HD mouse models. For example, an increased activity measured by c-Fos expression level was reported in the DMS but not in the DLS of R6/1 mice before motor deficit onset during an operant conditioning learning task (Cabanas et al., 2017). In addition, alterations of excitatory synaptic transmission from motor cortex to DLS of was reported in YAC128 mice, accompanied with behavioral deficits during a pellet reaching task (Glangetas et al., 2020).

While these studies suggest motor skill deficits in HD transgenic mice that over-express mutant HTT or fragments of mutant HTT at high level in a context of two wild-type alleles, our study was designed to explore whether deficits in motor learning occur early in the disease progression, in a mouse model of HD that recapitulates the genetic profile of HD patients and that develops motor phenotypes late during their lifespan (Hdh^CAG140/+^). Using the accelerating rotarod paradigm, we evaluated the motor learning performance of young Hdh^CAG140/+^ mice (1.5 month-old) and found a decreased performance in the late phase. We had shown previously that DMS and DLS are differentially involved in the progression of motor learning, with the formation of transitory DMS representations in the form of highly active cells in the initial training phase and long-lasting clusters of activity in DLS at late stages (Badreddine et al., 2022). In young Hdh^CAG140/+^ mice, *ex vivo* two-photon calcium recordings revealed alterations of basal network activity, in both the DMS and the DLS. This altered basal activity precluded the emergence of neuronal representations of initial and late motor learning after training observed in WT mice. Thus, our data show early deficits in motor learning, which were associated to dysfunctions in striatal networks. These alterations and deficits, occurring in an early phase before motor deficits occur, could be a promising way of identifying early markers of the disease.

## RESULTS

### Early phase of motor learning is not significantly affected in young Hdh^CAG140/+^ mice

We used the accelerating rotarod paradigm (Badreddine et al., 2022; Costa et al., 2004; Kupferschmidt et al., 2017) to evaluate motor learning in young (1.5 month-old) Hdh^CAG140/+^ mice and age-matching wild-type (WT) mice. We considered two distinct training stages, early and late phases, that we recently characterized with significant activity changes in DMS and DLS respectively (Badreddine et al., 2022). This allowed us to evaluate the possible progressive alterations of performance in Hdh^CAG140/+^ mice at different stages of motor learning, and in two functional territories of the dorsal striatum.

We first evaluated the performance of mice during the early training phase, i.e. corresponding to a single training session of 10 trials, with 5 minutes resting period between each trial (Fig.1a). We trained the two groups of mice, WT and Hdh^CAG140/+^, and compared their performance. We did not observe any marked difference in the latency-to-fall curve between the two groups after early training (p=0.1794, F_(1,28)_=1.896, two-way ANOVA) (Fig.1b). We used a quantitative measure with a one-phase association fit and the slopes of the fitted curves were not significantly different (p=0.5455, t-test) (Fig.1c). It should be noted that plotting the individual performance highlighted a higher inter-individual variability in the initial performance of Hdh^CAG140/+^ mice compared to WT mice (Fig. 1d). Nevertheless, both groups had a higher performance at the end of training compared to the beginning (WT mice: p<0.0001; Hdh^CAG140/+^ mice: p=0.0029, paired t-test) (Fig.1d). These results were confirmed by the learning index (LI), a quantification of the performance of the mice, calculated as the difference between the beginning and end of training (LI = last two trials – first two trials) (Fig.1e). When comparing the LI of WT and Hdh^CAG140/+^ mice, we did not see any significant difference between the two groups (unpaired t-test, p=0.7248). Therefore, there were no strong alterations of the early stage of motor learning in Hdh^CAG140/+^ mice, although the variability was higher compared to WT mice.

**Figure 1:**
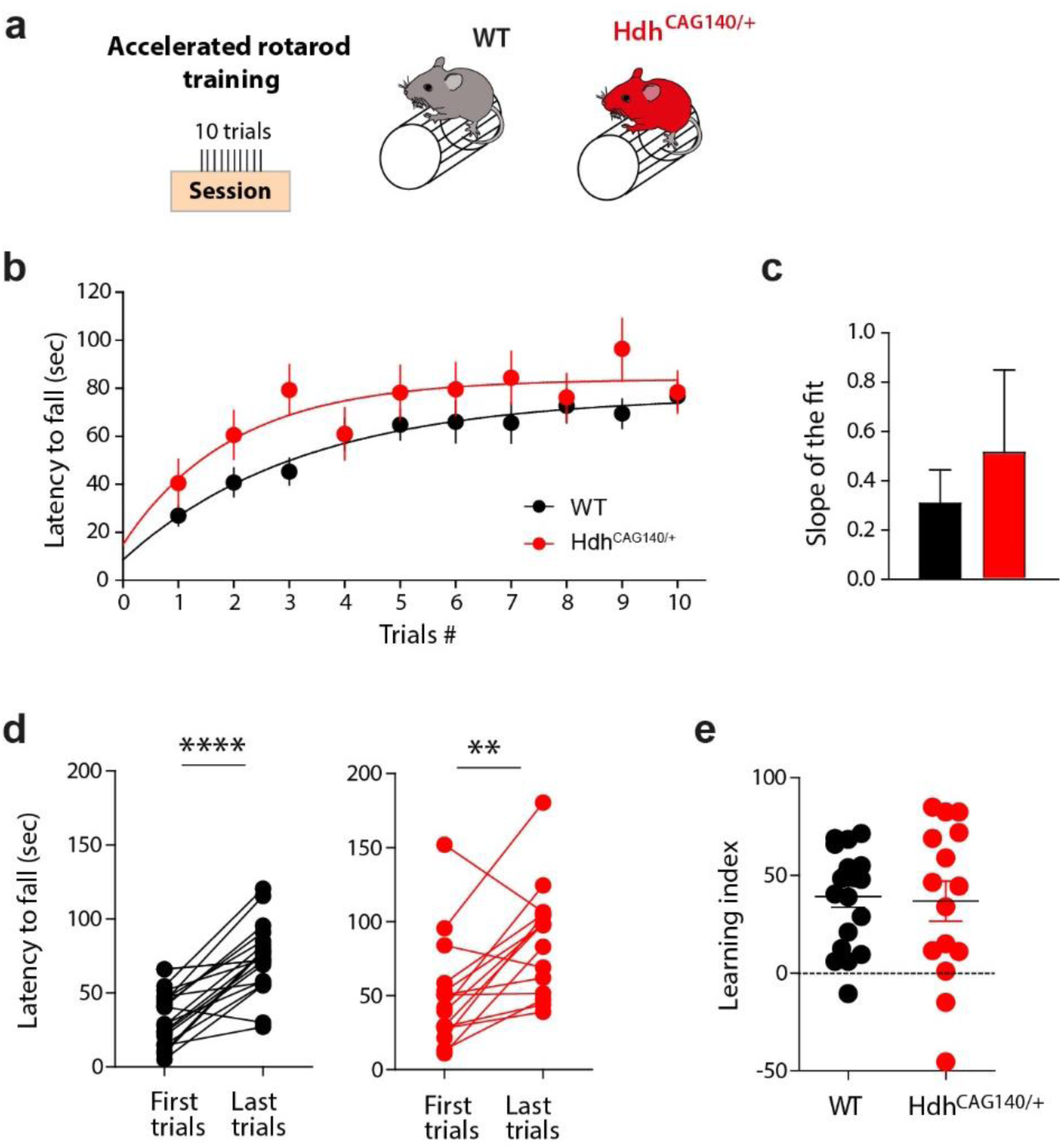
Similar behavior of WT and Hdh^CAG140/+^ mice during early phase of rotarod training. **a**: Behavioral paradigm used to study the early phase of motor learning: WT and Hdh^CAG140/+^ mice are trained on an accelerating rotarod for 1 session of 10 trials (early training). **b**: learning curve of WT (black, n=15) vs. Hdh^CAG140/+^ (red, n=15) mice. No difference in performance was found between both groups of mice (p=0.1794, F_(1,28)_=1.896, two-way ANOVA). **c**: Slopes of the one-phase association fits of the learning curve. No significant difference was observed between both groups (p=0.5455, t-test). **d**: Latency to fall during the first and last trials of Day1. Both groups increase their performance between the beginning and the end of training (WT: p<0.0001, Hdh^CAG140/+^: p=0.0029, paired t-test), with more variability in the Hdh^CAG140/+^ group. **e**: Learning index was not significantly different between WT and Hdh^CAG140/+^ mice (p=0.7248, unpaired t-test).

### DMS network dynamics are altered in young Hdh^CAG140/+^ mice

Even though we did not observe any major deficits in early behavioral performance of Hdh^CAG140/+^ mice, we explored whether there were any alterations in the DMS dynamics. To do so, we injected AAV-GCAMP6f in the DMS and probed DMS network activity with *ex vivo* two-photon calcium imaging in naïve and early-trained groups of WT and Hdh^CAG140/+^ mice (Fig.2a). We analyzed cortically-evoked activity in striatal projection neurons (SPNs) in parasagittal brain slices preserving cortical cingulate afferents to DMS (Badreddine et al., 2022; Fino et al., 2018). The amplitude of responses was measured for each cell and used to build DMS functional maps (Fig.2b). We first assessed whether the basal DMS activity was altered in the HD mouse model. We measured the mean amplitude of all SPN stimulation-evoked responses in naïve WT and Hdh^CAG140/+^ mice and observed that it was significantly lower in Hdh^CAG140/+^ animals, (n= 7 slices, n= 5 mice) compared to WT mice (n=10 slices, n= 10 mice) (p=0.0006, t-test) (Fig.2c). We searched for specific distribution of DMS activity by evaluating the percentage of highly active cells (HA cells), i.e. the percentage of cells with amplitude responses superior to the averaged amplitude of naive animals. The percentage of HA cells was similar between the two mice groups, with about half of the SPNs being active (WT vs Hdh^CAG140/+^) (p=0.9570, t-test) (Fig.2c). This means that even though Hdh^CAG140/+^ mice start with a lower amplitude of response, they exhibit the same pattern of activity than WT mice. Overall, the basal DMS activity is lower in Hdh^CAG140/+^ mice.

During initial phases of associative learning or motor learning, DMS neural activity displays an overall transitory decrease (Badreddine et al., 2022; Cataldi et al., 2022; Vandaele et al., 2019). In addition, we showed in previous work that such early DMS activity decrease unveils the emergence of a sparse set of HA cells, whose number is directly linked to the level of performance (Badreddine et al., 2022). We thus next explored whether network activity remapping associated to initial motor training was affected in Hdh^CAG140/+^ mice. For this purpose, we compared the DMS amplitude and activity patterns between naive and early-trained animals, in WT and Hdh^CAG140/+^. In order to better characterize any alterations that might occur in Hdh^CAG140/+^ mice, we included in the analysis all the animals that have followed early training, whatever the level of performance. In WT mice, we observed a decrease in SPNs overall activity in early-trained mice (n= 11 slices, n= 10 mice) compared to naive animals (Naive, n= 10 slices, n= 10 mice) (p=0.0044, t-test) (Fig.2d). On the contrary, in Hdh^CAG140/+^ mice we did not observe any difference in overall activity amplitude between naïve and early-training conditions (n= 7 slices, n= 5 mice) (p=0.3519, t-test) (Fig. 2f). The training thus did not lead to the same decrease of the DMS level of activity in Hdh^CAG140/+^ mice. The sparsity of highly active SPNs throughout the field are tightly correlated with early performance of motor learning (Badreddine et al., 2022). In WT mice, we confirmed that the percentage of HA cells significantly decreased in early-trained compared to naive mice (Naive, 47%, n= 10 slices, Early, 25%, n= 11 slices, p=0.0046, t-test) (Fig.2d). In addition, now considering the level of performance of the animals (i.e. the learning index), we observed that the proportion of HA cells was directly correlated to the animal’s performance (r²=0.5189, p=0.0124) (Fig.2e) in WT mice. We asked whether this was the case in the DMS activity of Hdh^CAG140/+^ mice after training. The proportion of HA cells did not change between naïve and early-trained animals (Naïve, 47%, n= 7 slices, Early, 50%, n= 7 slices, p=0.8353, t-test) (Fig.2f). Strikingly, in Hdh^CAG140/+^ early-trained mice, there was no correlation between HA cells and mice performance (r²=0.01816, p=0.7733) (Fig. 2g). Therefore, in Hdh^CAG140/+^ animals, the basal activity is altered, with overall lower activation of DMS SPNs. This precluded any reorganization of DMS activity associated with early motor training and any link with mice performance.

**Figure 2:**
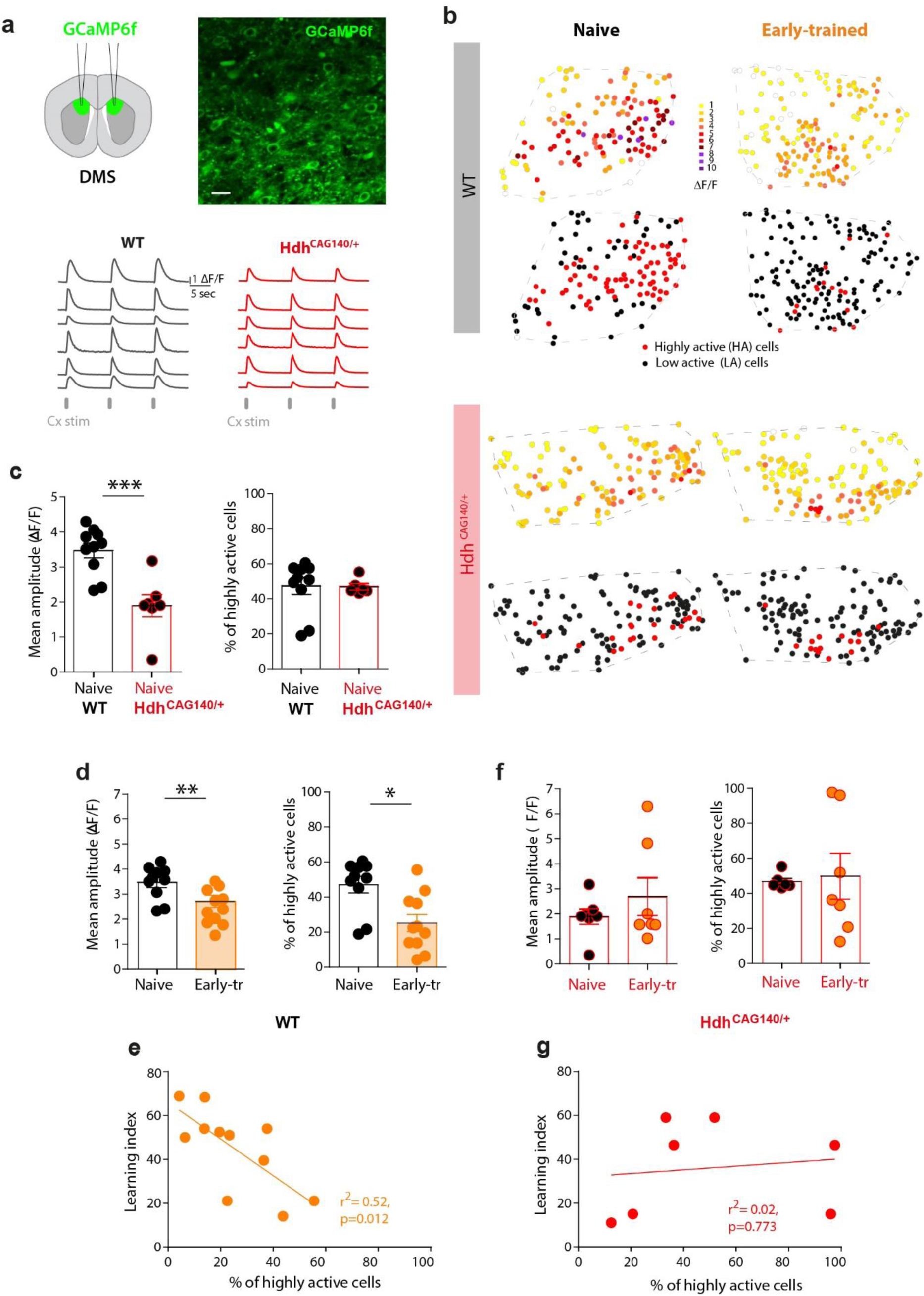
Basal DMS activity and early-training induced plasticity are altered in young Hdh^CAG140/+^ mice. **a**: Experimental conditions: injections of calcium sensor GCaMP6f in the DMS. Two-photon calcium imaging acquisition of a field of view showing SPNs expressing GCaMP6 in the DMS (scale bar: 20µm) and representative calcium transients evoked by cortical stimulations in either WT or Hdh^CAG140/+^ mice. **b**: Representative functional maps of striatal networks in DMS for naïve and early-trained conditions, in WT and Hdh^CAG140/+^ mice. The color code corresponds to the amplitude of response (ΔF/F). Bottom maps present the HA (highly active) cells in red and low active (LA) cells in black. **c**: Top graph: Averaged amplitude of response of all SPNs in the recording field in naïve animals (WT n=10, Hdh^CAG140/+^ n=7, p=0.006, t-test). Bottom: Percentage of HA cells in naïve animals of both groups (p=0.9570, t-test). **d**: Mean amplitude of response and % of HA cells in naïve (black) and early-trained (orange) WT mice. Early training leads to a significant decrease of overall activity (p=0.0044, t-test), and of the % of HA cells (p=0.0046, t-test) compared to naïve conditions in WT mice. **e**: Correlation between the percentage of HA cells and the learning index of the animals after early training. Significant correlation for the WT early-trained mice in DMS (r²=0.5189, p=0.0124, Pearson correlation). **f**: No difference neither in mean amplitude (p=0.3519, t-test), nor in % of HA cells (p=0.8353, t-test) between naïve and early-trained Hdh^CAG140/+^ mice. **g**: No significant correlation between the percentage of HA cells and the learning index of the mice after early training for Hdh^CAG140/+^ early-trained mice (r²=0.01816 p=0.773, Pearson correlation).

Overall, we observe alterations of the dynamics of the DMS networks without significant alterations of early training in Hdh^CAG140/+^ mice. This suggests that such network alterations may not be strong enough to exert a noticeable influence on behavior during the initial training phase but might affect a proper development of motor learning at later stages.

### Late stage of motor learning is affected in young Hdh^CAG140/+^ mice, with no locomotor or motor coordination deficits

We next asked whether the performance of Hdh^CAG140/+^ mice was affected after a long motor training. The mice were tested with a course of 7 consecutive days of rotarod training, one session a day, with 10 trials per session (Badreddine et al., 2022; Jin & Costa, 2015; H. H. Yin et al., 2009). We trained both groups of mice with the full training protocol (Fig.3a) and observed that Hdh^CAG140/+^ mice had altered performance compared to WT mice, with a significantly different latency-to-fall curve (p<0.0001, F_(2,2170)_=2.50, two-way ANOVA). The quantitative measure using one-phase association fit showed a significant difference in the plateau of the curves, which was significantly lower in Hdh^CAG140/+^ mice (p<0.0001, t-test) (Fig.3b). These results were more striking when considering the individual mouse performance in each group and comparing first and last trials. Even though both WT and Hdh^CAG140/+^ mice were able to learn the task (higher performance on the last day compared to the first one), Hdh^CAG140/+^ mice had a lower progression compared to WT (Fig.2c). Indeed, as a quantification, the learning index (LI) of Hdh^CAG140/+^ mice was significantly lower compared to WT mice (p=0.0322, t-test) (Fig.3 c and d). Altogether we observed that motor training was affected in Hdh^CAG140/+^ mice compared to WT mice.

**Figure 3:**
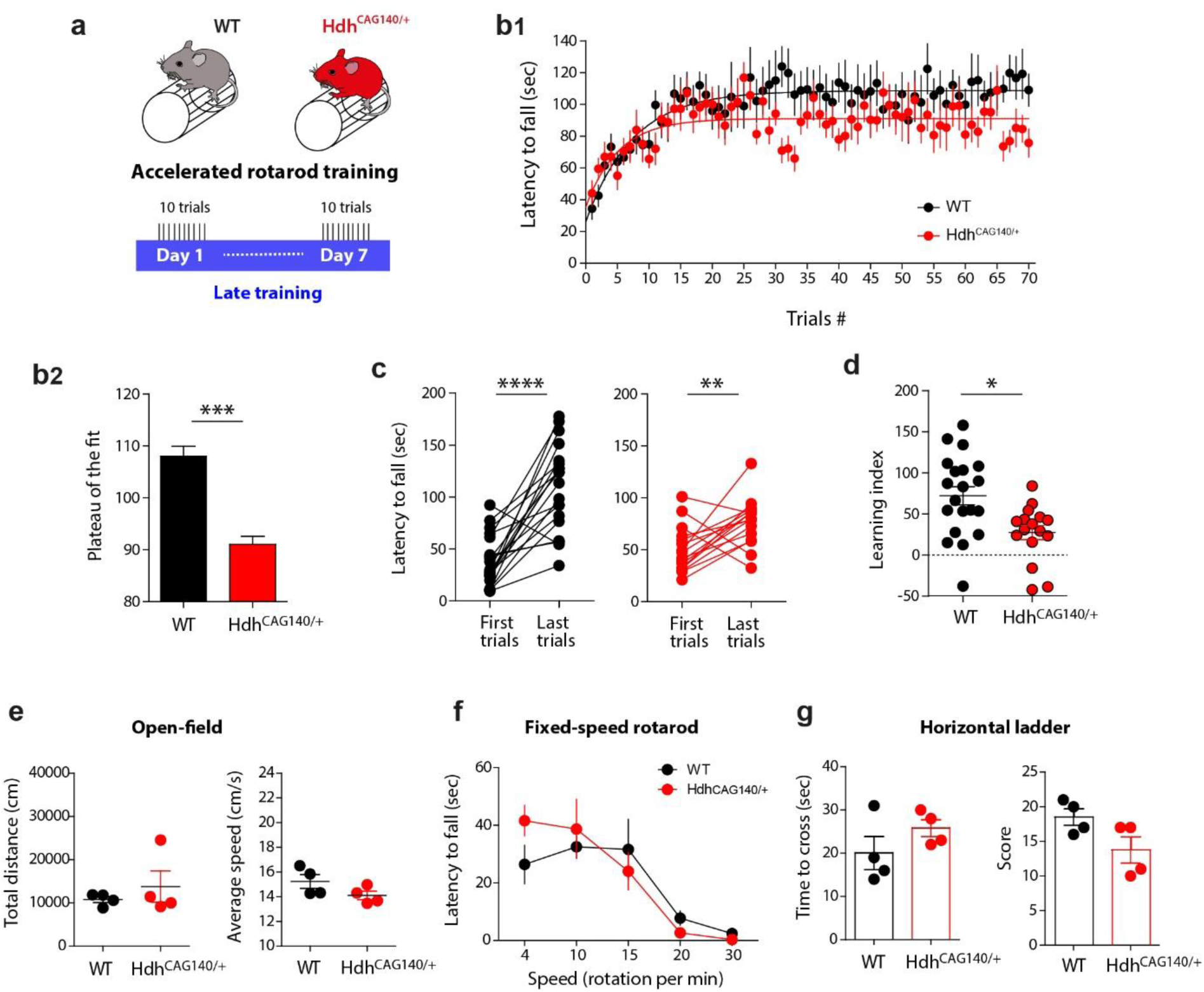
Deficits in late phase of motor learning but not in locomotor activity and motor coordination in young Hdh^CAG140/+^ mice. **a**: Behavior paradigm used to perform motor learning: WT and Hdh^CAG140/+^ mice were trained on an accelerating rotarod for 7 days, 1 session per day, 10 trials per session. **b**: **b1**: learning curves and one-phase association fits of WT (n=21 mice) vs Hdh^CAG140/+^ (n=17 mice) mice performance. Hdh^CAG140/+^ mice have a significantly lower performance compared to WT mice (p<0.0001, F_(2,170)_=2.503, two-way ANOVA). **b2**: Significant difference between plateau of WT and Hdh^CAG140/+^ mice (p< 0.0001, t-test). **c**: Latency to fall of the first trials of day1 and the last day of training (Day 7). Both groups increase their performance between the beginning and the end of training (WT: p<0.0001, Hdh^CAG140/+^: p=0.0011, paired t-test), with lower performance for Hdh^CAG140/+^ mice. **d**: Learning index of Hdh^CAG140/+^ mice is significantly lower compared to WT mice (p=0.0322, t-test). **e**: Open-field test. Total distance travelled (p=0.4465, t-test) and average speed of locomotion (p=0.1341, t-test) were not significantly different between WT (n=4) and Hdh^CAG140/+^ (n=4) mice. **f**: Fixed-speed rotarod. Four different speeds, 4, 10, 20 and 30 rpm, were tested. No significant difference was observed between WT and Hdh^CAG140/+^ mice (p=0.7288, F_(1,30)_=0.1224, two-way ANOVA). **g**: Horizontal ladder test: the time to cross the ladder and the score were measured. No significant difference between WT and Hdh^CAG140/+^ mice neither for time to cross the ladder (p=0.2267, t-test), nor for the score (p=0.0773, t-test).

We next wanted to ensure that this alteration of the performance of young Hdh^CAG140/+^ mice was not due to locomotor activity or motor coordination deficits. In a subset of mice, we assessed motor behavior with different tests, open-field, fixed-speed rotarod and horizontal ladder. Global locomotor activity was tested in an open-field: the travelled distance and the averaged speed were comparable between Hdh^CAG140/+^ (n=4 mice) and WT mice (n=4 mice) (distance: p=0.4465, speed: p=0.1341, t-test) (Fig.3e). Motor coordination was first assessed with the fixed-speed rotarod paradigm and showed that Hdh^CAG140/+^ mice remained on the rod for the same amount of time than WT mice at the different measured speeds (p=0.7288, F_(1,30)_=0.1224, two-way ANOVA) (Fig.3f). For a finer motor coordination assessment, we used the horizontal ladder test. Again, Hdh^CAG140/+^ mice and WT mice spent the same amount of time to cross the ladder (p=0.2267, t-test), and had a similar score (p=0.0773, t-test) (Fig.3g). Our results concord with the literature characterizing the Hdh^CAG140/+^ mouse model that does not show any motor deficits in the an early phase (Menalled et al., 2003), the same phase we examined in this study (1.5 month-old mice). We thus show that the deficits in motor learning are not due to major motor deficits from the development of the disease.

### Motor learning deficits are associated with DLS network alterations

We next questioned whether the late training deficits of Hdh^CAG140/+^ mice were associated with altered DLS network activity, using *ex vivo* DLS calcium imaging in 1.5 month-old mice (Fig. 4a). We measured the averaged cortically-induced activity per DLS field, in the somatosensory (S2) projecting area (Fig. 4b). Comparing SPNs overall activity in naïve WT and Hdh^CAG140/+^ mice, we observed a significantly lower mean amplitude in Hdh^CAG140/+^ mice (n= 11 slices, n= 9 mice) compared to the WT (n= 9 slices, n= 9 mice) (p=0.002, t-test) (Fig. 4c). Therefore, like in DMS, Hdh^CAG140/+^ mice had a lower basal activity level in DLS compared to WT mice. We also explored the DLS activity patterns in WT and Hdh^CAG140/+^ mice by measuring the area formed by the cells with the highest activity (HA area). We observed a significantly smaller HA area in Hdh^CAG140/+^ naïve mice compared to the WT group, consistent with the overall lower level of activity (p=0.0011, t-test) (Fig. 4c). Thus, naïve Hdh^CAG140/+^ mice have a lower and smaller active area within the DLS compared to WT mice. We next evaluated whether Hdh^CAG140/+^ mice had any deficits in network’s plasticity. To test this, we built functional maps of the same field of view at two different frequencies of cortical stimulation (5 and 20 Hz) and extracted the mean SPN amplitude and HA area. WT naïve mice displayed plastic properties, i.e. adaptation of activity levels to increased cortical inputs stimulations from 5 to 20 Hz. We observed an increase in the mean amplitude of response (p=0.0003, paired t-test), but also a widening of the HA area (p=0.0081, paired t-test) when increasing cortical stimulations, consistent with previous results (Badreddine et al., 2022). For Hdh^CAG140/+^ mice, although starting from a lower level, there was also an increase of the mean activity with increasing frequencies (Fig.4d) (p=0.0003, t-test). Nevertheless, the HA area did not significantly change with increasing frequency stimulations (p=0.6696, paired t-test). Overall, these results show that DLS basal activity and network plasticity are strongly affected in young Hdh^CAG140/+^ mice.

**Figure 4:**
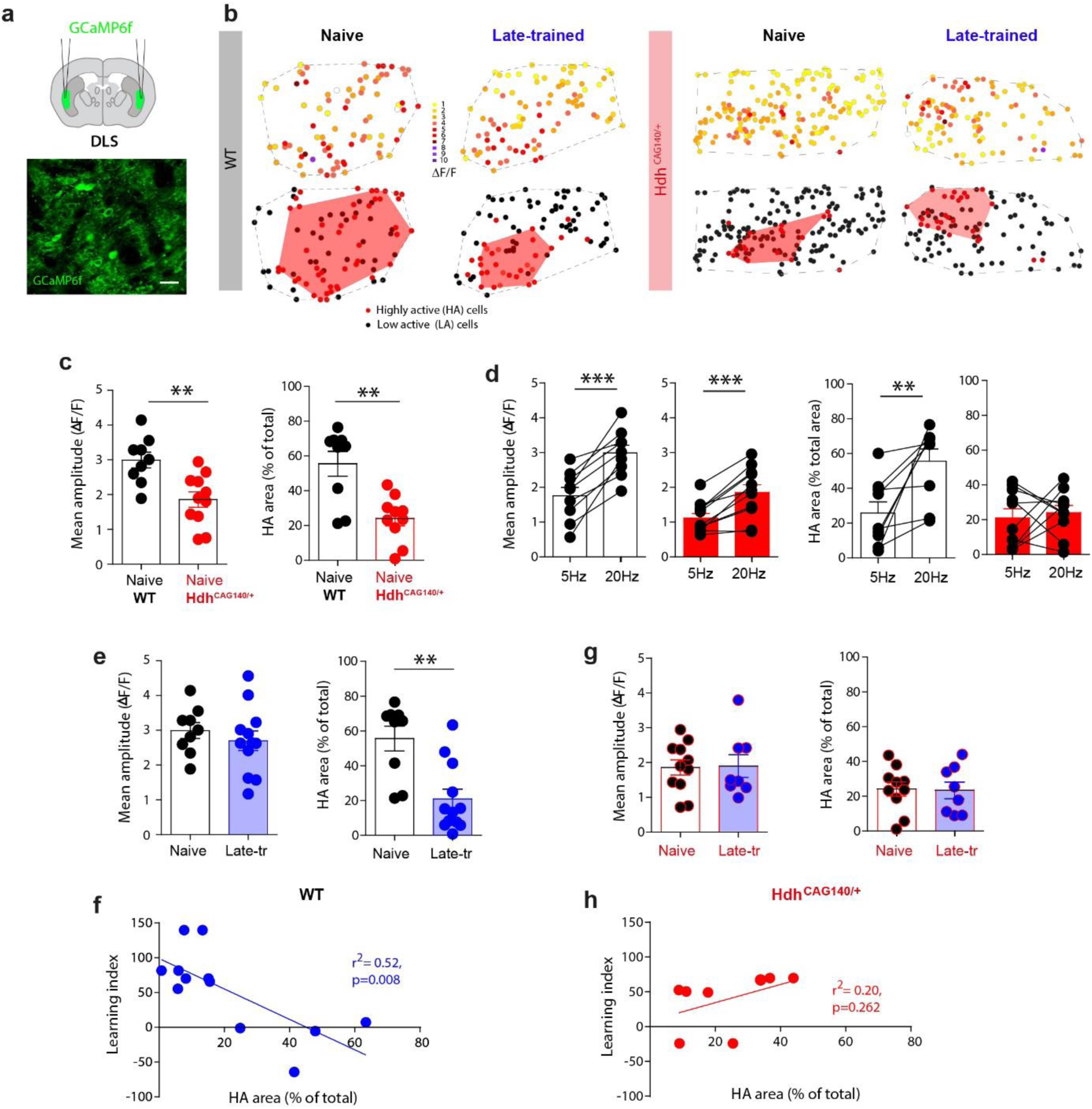
Alterations of basal DLS activity and late-training induced plasticity in young Hdh^CAG140/+^ mice. **a**: Experimental conditions: injection of calcium sensor GCaMP6f in the DLS in WT and Hdh^CAG140/+^ mice and two-photon calcium imaging acquisition of a field of view showing SPNs expressing GCaMP6 in the DLS (scale bar: 20µm). **b**: Representative functional maps of striatal networks in DLS for naïve and late-trained conditions, in WT and Hdh^CAG140/+^ animals. The color code corresponds to the amplitude of response (ΔF/F). Bottom maps present the HA cells in red and LA cells in black. **c**: In naïve animals, the averaged amplitude of response of all SPNs in the recording field is lower in Hdh^CAG140/+^ mice (WT, n=9 slices from 9 mice, vs Hdh^CAG140/+^, n= 11 slices from 9 mice, p=0.0022, t-test). Similarly, the percentage of HA area, the area of the field corresponding to clustered activity, is also significantly lower (p=0.0011, t-test). **d**: Plasticity properties of the network was assessed by measuring the striatal activity adaptation to increasing stimulation frequencies of cortical inputs. The mean amplitude increases between t 5 and 20Hz for both WT (left, p=0.0003, paired t-test) and Hdh^CAG140/+^ mice (right, p=0.0003, paired t-test). The percentage of HA area also increases between 5 and 20Hz for WT mice (p=0.0053). On the contrary, Hdh^CAG140/+^ mice displayed no significant change in HA area in response to increasing frequencies (p=0.6696). **e**: Averaged amplitude of response in WT mice, in naïve and late-trained conditions (n= 12 slices, n=11 mice). We observed no significant difference in amplitude between naïve and late-trained, but a decrease in HA area, indicating a clustering of the DLS activity (amplitude: p=0.4553, HA area: p=0.0031, t-test). **f**: Correlation between the HA area and the learning index of the animals after late training. Significant correlation for the WT late-trained mice in DLS (r²=0.5181, p=0.0083). **g**: Averaged amplitude of response in Hdh^CAG140/+^ mice, in naïve and late-trained conditions (n= 8 slices, n= 7 mice). There was no significant difference in amplitude, nor in HA area (amplitude: p=0.9170, HA area: p=0.9072, t-test). **h**: No significant correlation between learning index and % of HA area in Hdh^CAG140/+^ late-trained mice (r²=0.2035, p=0.2619).

We next investigated whether motor training-induced spatiotemporal re-organization of DLS activity was affected in Hdh^CAG140/+^ mice after late training, including all trained animals. First, measuring the overall DLS SPN amplitude, we observed no difference in the late-trained WT mice (n = 12 slices, n= 11 mice) compared to the naïve WT mice (p=0.4553, t-test) (Fig. 4e). This was similar in Hdh^CAG140/+^ mice, for which late rotarod training had no effect on the amplitude of responses (Hdh^CAG140/+^ n= 8 slices, n = 7 mice, p=0.9170, t-test) (Fig. 4g).

We previously described that the DLS activity is spatially restricted in clusters after late motor learning (Badreddine et al., 2022). Indeed, in WT mice, there was a significant decrease of active area (HA area) after late training (p=0.0031, t-test) (Fig. 4e). In addition, considering the level of performance of the animals, the DLS active area was significantly correlated with the learning index (r²=0.5181, p=0.0083); i.e. the smaller the clustering area, the better the animal learns (Fig. 4f). We asked whether such training-associated DLS activity re-organization was affected in Hdh^CAG140/+^ mice. We observed that in Hdh^CAG140/+^ mice, the HA area was not modified by late training (p=0.9072, t-test) (Fig.4g), and was not correlated with the motor performance (r²=0.2035, p=0.2619) (Fig.4h). There was therefore no learning-associated activity remapping of DLS in the young Hdh^CAG140/+^ mice. This was probably due to an altered DLS activity in the basal conditions in naïve Hdh^CAG140/+^ mice.

Altogether these results show that DLS networks are altered in Hdh^CAG140/+^ mice in a naïve state and these basal changes strongly affect the DLS dynamics and the performance in motor learning, without major alterations in motor abilities.

### DLS network alterations are not modified by the appearance of motor deficits in older _HdhCAG140/+ mice_

Mice used in our study so far were young (1.5 month-old) and did not present deficits in motor activity and coordination (Fig.3). Nevertheless, we observed alterations in DLS activity dynamics and motor learning at that stage. We thus wondered whether the occurrence of deficits in motor coordination, later in the lifespan of Hdh^CAG140/+^ mice, might affect the DLS activity networks. In order to answer this question, we used 5 month-old Hdh^CAG140/+^ mice, when motor deficits have been reported (Menalled et al., 2003). We first confirmed in our conditions whether Hdh^CAG140/+^ mice at this age exhibited signs of impaired motor activity/coordination, compared to age-matching WT mice. We observed no deficit in locomotor activity in the open-field, as there was no difference between WT (n= 12 mice) and Hdh^CAG140/+^ (n= 9) mice, (distance travelled: p=0.4823, average speed: p=0.9299, t-test) (Fig. 5c). We next used finer tests to assess motor coordination. On the fixed-speed rotarod, we observed a lower latency to fall in Hdh^CAG140/+^ compared to WT mice (p=0.0377, F_(1,34)_=4.678, two-way ANOVA) (Fig. 5a). These results were confirmed with the horizontal ladder test since we reported a significantly lower score and time to cross the ladder for Hdh^CAG140/+^ mice (time to cross: p=0.0081, score: p=0.0210, t-test) (Fig.5b). There are thus deficits in motor coordination of 5 month-old Hdh^CAG140/+^ mice.

**Figure 5:**
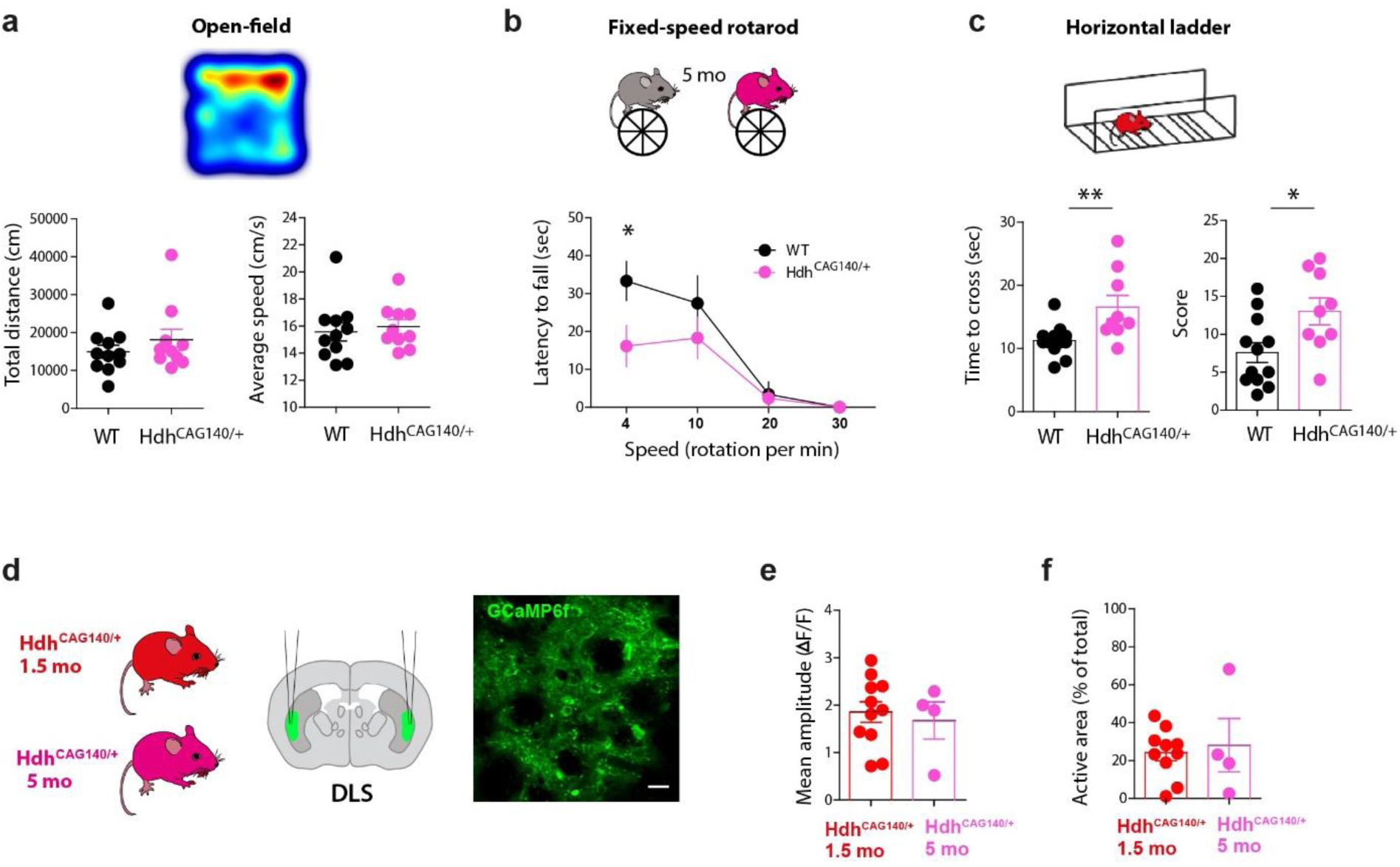
Apparition of motor deficits in symptomatic Hdh^CAG140/+^ mice does not change the alterations in DLS activity. **a**: Open-field test. Total distance travelled (p=0.4823, t-test) and average speed of locomotion (p=0.9299, t-test) were not significantly different between WT (n= 12) and Hdh^CAG140/+^ (n=9) mice. **b**: Fixed-speed rotarod with four different speeds tested. Hdh^CAG140/+^ mice had a lower performance on the fixed-speed rotarod compared to WT mice (p=0.0377, F_(1,34)_=4.678, two-way ANOVA). **c**: Horizontal ladder test. Hdh^CAG140/+^ mice spent more time to cross the ladder than WT mice (p=0.0081, t-test), and had a higher score (p=0.0210, t-test). **d**: Injection of GCaMP6f in the DLS of 5 months-old mice and comparison with similar injections in younger (1.5 month-old) Hdh^CAG140/+^ mice. **e**: Measure of DLS activity in 1.5mo (n= 10) and 5mo (n= 4) Hdh^CAG140/+^ mice. No significantly different amplitudes observed between 1.5 and 5 months-old mice (p=0.6846, t-test). **f**: The pattern of activity measured with the cluster area (HA area) was not significantly different (p=0.7159, t-test).

We tested whether motor impairments would significantly affect striatal activity patterns. Since the DLS displayed sufficient alterations that might explain the motor learning impairment, we focused on the effect of motor coordination deficits on the DLS network’s activity, using ex vivo calcium imaging in 5 month-old Hdh^CAG140/+^ animals (n=4 mice). We performed similar analysis as before by first measuring the level of overall activity and then examining the potential clustering of the activity and comparing it with young (1.5 month-old) Hdh^CAG140/+^ mice. We observed no difference in the mean activity between naïve young and older Hdh^CAG140/+^ mice (p=0.6846, t-test) (Fig.4e). Similarly, the clustering of highest activity was not different between naïve young and older animals (p=0.7159, t-test) (Fig.4f).

Altogether, these results suggest that the alterations of DLS networks, by not following the same reorganization as in WT mice after training, might be responsible for the deficits in motor learning. And these alterations are not affected by the occurrence of motor deficits at later stages of HD. Thus, the alterations occurring in the dorsal striatum might be a good indicator for the motor learning deficits observed in HD.

## DISCUSSION

In this study, we showed strong deficits in motor learning in young animals in a mouse model of HD (Hdh^CAG140/+^), when no locomotor activity nor motor coordination impairments manifested yet. These alterations are particularly important in the late consolidation phase of the training. Such behavioral affection is linked to striatal network dysfunctions. Indeed, Hdh^CAG140/+^ mice displayed lower activity both in the DMS and the DLS in basal conditions compared to WT animals. These basal differences precluded the spatiotemporal re-organization of striatal activity associated with different stages of motor learning.

### Motor learning alterations and associated striatal activity dysfunctions in young Hdh^CAG140/+^ mice

We show here that alterations of motor learning in 1.5 month-old Hdh^CAG140/+^ mice were not due to locomotion or motor deficits. This is consistent with report in the Hdh^CAG140/+^ mice of motor symptoms developing in older animals, after 4 months of age (Hickey et al., 2008), which we showed previously in 7 month-old mice (Virlogeux et al., 2021), and here in 5 month-old mice (Fig.5). Another study using the same Hdh^CAG140/+^ mouse model, showed that deficits on the accelerated rotarod did not appear before 11 months (Rising et al., 2011). This discrepancy probably stems from the different training protocols, as in this study the accelerated rotarod paradigm was done over a much shorter period of 3 days with 4 trials per session.

In order to link behavior alterations and striatal activity, we explored the activity patterns of both DMS and DLS networks, as they are differentially recruited throughout motor learning. In Hdh^CAG140/+^ mice, we showed that DMS networks were altered in a basal condition, before training, and did not exhibit any reorganization upon training. These alterations might not have been sufficient to affect significantly the early training aspect of the behavior but showed a higher variability in Hdh^CAG140/+^ mice performance and could contribute to defects in the late training phase. We indeed reported that motor learning was particularly affected in the late phase in Hdh^CAG140/+^ mice as they reached a lower plateau of the learning curve compared to WT mice. The deficits in the performance of Hdh^CAG140/+^ mice on the accelerated rotarod were here reflected by alterations in the DLS networks activity patterns; the overall DLS activity was lower with a more restricted active area in naïve Hdh^CAG140/+^ mice, which was not changed by full training. Interestingly, the active cluster area was smaller compared to naïve WT mice and bigger compared to late-trained WT mice. These results suggest not only a dysfunction in the DLS networks in basal conditions, but also highlight the lack of plasticity of the DLS networks and the absence of formation of smaller clusters after training. This disorder in plasticity is really interesting and should be a promising venue to understand network dysfunctions in HD models.

### Mechanisms of striatal networks alterations in HD

Understanding the mechanisms responsible for functional alterations is key to identify eventual targets to rescue the dysfunctions. Alterations can appear in cellular physiology, synaptic transmission or even in anatomical connectivity. We previous reported differential mechanisms associated with motor learning on the DMS and DLS networks plasticity: synaptic plasticity of cortical inputs in DMS, and anatomical plasticity in DLS with an increase in the number of somatosensory inputs into DLS after late training (Badreddine et al., 2022). In HD rodent models, altered synchrony has been described between cortical and striatal networks (Lee Hong & Rebec, 2012; Naze et al., 2018). In addition, Deng et al. showed a decrease of the number of corticostriatal terminals in the DLS in 12 month-old Hdh^CAG140/+^ mice (Deng et al., 2013). Recently a reduced functional connectivity between different cortical areas and the DLS was described in a symptomatic R6/1 mouse model of HD (Conde-Berriozabal et al., 2024). An elevated cortical activity has also been described early in the disease in animal models with imaging and electrophysiological studies (Arnoux et al., 2018; Burgold et al., 2019; Donzis et al., 2020). Based on these data, we could expect different mechanisms for the Hdh^CAG140/+^ mice compared to WT mice. Indeed, these results suggest alterations in the corticostriatal projections and a poorer synaptic plasticity. Our lab recently showed that a phosphorylation of the huntingtin protein leads to an over-accumulation of synaptic vesicles at the synapse and an increased probability of release from M1 to DLS which leads to altered motor learning (Vitet et al., 2023). Interestingly, a recent study using optogenetics showed that a repetitive stimulation of the M2 to DLS projections not only reversed motor deficits in 12 month-old mice, but also increased synaptic plasticity (Fernández-García et al., 2020). We can thus wonder if the mechanism would be similar in our conditions of somatosensory to DLS projections and if repetitive training, stimulating those S2-DLS projections, might be able to restore the observed deficits in Hdh^CAG140/+^ mice. Synaptic plasticity from the M2 onto the DLS is reduced in HD mouse models (Fernández-García et al., 2020; Glangetas et al., 2020; Hintiryan et al., 2016; Raymond et al., 2011). We explored the plasticity of the corticostriatal networks by examining the response at low and high frequencies (5 and 20Hz). In the DLS, we noted an absence of plasticity where we observed similar cluster areas at low and high stimulation frequencies, suggesting the corticostriatal dysfunctions might be responsible for the deficits in late learning. Our data corroborate the report of a reduction in the projections from the cortex to the striatum that only concerned M2 to DLS projections (Hintiryan et al., 2016) and more recent studies reporting this decrease between different cortical areas and specifically the DLS but not the DMS (Conde-Berriozabal et al., 2024). Previous studies focused mainly on the M2 to DLS projections because of the characteristic motor deficits in HD. However, with new evidence of alterations between DLS and other cortical structures, future work should explore more broadly the different mechanisms behind the alterations in motor learning and the differential alterations of functional striatal territories.

### Alterations of striatal activity and motor learning as early markers of the disease?

Our study shows an altered performance in young Hdh^CAG140/+^ mice, without any major motor deficits. In 5 month-old Hdh^CAG140/+^ mice, motor symptoms manifested with altered motor coordination and fine motor control. Since the global DLS network properties were not affected by motor coordination deficits in older animals, it could suggest a specificity of these alterations to motor learning independently of the later motor symptoms. Future work should extend this observation to other learning contexts using other behavioral tests, to confirm our observations. Indeed, if confirmed, the alteration of motor learning could be a potent early marker of the disease onset. One of the directions for HD research this past decade has been to find markers for the onset of the disease. The PREDICT-HD study measured the performance of asymptomatic HD patients with a confirmed CAG expansion on 21 standardized cognitive tasks (Stout et al., 2011), with motor learning testing as one of these tasks. This type of early detection could allow adjusting potential therapeutic targets for the patients.

## MATERIALS AND METHODS

### EXPERIMENTAL MODELS

Hdh^CAG140/+^ knock-in mice with a C57BL/6J background were used in this study. This mouse model was chosen because Hdh^CAG140/+^ mice develop motor symptoms after several months (Crook & Housman, 2011; Menalled et al., 2003), allowing us to study the early phase preceding any motor deficits. Moreover, these mice express human HTT exon1 with 140 repeats of CAG. Mice used in this study were all heterozygous to replicate the HD in humans. Mice were 1.5 or 5 months old. They were housed in temperature-controlled rooms with standard 12 hours light/dark cycles and food and water were available *ad libitum*. Every precaution was taken to minimize stress and the number of animals used in each series of experiments. All experiments were performed in accordance with EU guidelines (directive 86/609/EEC) and in accordance with French national institutional animal care guidelines (protocol APAFIS#29200-2016092317163976 v6).

### METHOD DETAILS

#### AAVs

An adeno-associated virus (AAV) serotype 5 was used to express a calcium indicator in striatal cells. AAV5-syn-GCaMP6f-WPRE-SV40 was purchased from UPenn Core (PA, USA) and Addgene.

#### Stereotaxic injections

Stereotaxic intracranial injections were used to deliver the AAV in the striatum. Mice were anesthetized with 2.5 % isoflurane and placed in a stereotaxic frame (Kopf). Under aseptic conditions, the skull was exposed and levelled, and a craniotomy was made with an electric drill. The viruses (serotype 5, ≈ 10^12^ genomic copies per mL) were injected through a pulled glass pipette (pulled with a P-97 model Sutter Instrument Co. pipette puller) using a nanoinjector (World Precision Instruments, Germany). The pulled glass micropipette was slowly lowered into the brain and left 1 min in place before starting the injection of the virus at an injection rate of 100 nL per min. A volume of 400 nL of the virus was enough to infect a large proportion of DMS or DLS. The injections targeted the DMS at coordinates AP + 1.2mm, ML 1.2, DV - 1.9 and the DLS at AP - 0.38, ML 2.3, DV - 2.45. Following injections, we waited 5 min before raising the pipette out of the brain. To minimize dehydration during surgery mice received a subcutaneous injection of 1mL of sterile saline. Post-operatively mice were kept on a heating pad for 1 h before being returned to their home cage. Mice were then checked daily for 4-5 days. A period of 15 to 20 days after injections was enough to allow for a good expression of AAVs. We observed similar expression of GCaMP6f in all striatal neurons in DMS or DLS in injected mice with viral vectors.

#### Behavioral training

##### Accelerating rotarod

In the previous days of the training, mice were habituated to the room and to handling. To assess motor learning, mice learned to run on a rotating rod (Panlab). For each trial the mouse was placed on the still rotarod which was activated at that point. The rotation of the rod was increasing from 4 to 40 rotations per min over 300 s. Each trial was ended when the mouse falls off the rotarod or when the 300 s had elapsed. There was a resting period of 300 s between trials. Animals were trained with 10 trials per day for either 1 day (early training) or 7 days every day (late training). This training protocol was chosen since it was previously described as a reliable test for motor skill learning or procedural learning.

##### Fixed-speed rotarod

To test motor balance and coordination of animals, a fixed-speed rotarod was used. The same apparatus was used (Panlab) with four fixed different speeds tested: 4, 10, 20 and 30 rpm. Animals were placed on the rotating wheel and their latency to fall was measured. Five trials per speed were done, with a 300 s resting period between each trial. A resting period of 600 s was given to mice before changing the speed of the rod.

##### Open-field

The open-field test was performed during the light phase of the light cycle to test locomotor activity. Mice were placed in the center of a square Plexiglas open-field (50×50×50 cm) for 60 minutes exploratory spontaneous motor activity session, while they were video-recorded with the Viewpoint VideoTrack system. Before each mouse was tested, the open-field was cleaned with ethanol.

##### Horizontal ladder

We used the horizontal ladder test to finely assess motor coordination. Two clear Plexiglas walls (69.5 x 15 cm) were linked by metal rungs (0.2cm in diameter) and elevated by 30 cm. The width of the alley was adjusted to the size of the animal to prevent the animal from turning around. Mice were habituated during two consecutive days (three trials per day) to walk on the horizontal ladder with regularly spaced rungs (2 cm apart). On the third day, mice were video-recorded while being tested on irregularly spaced rungs (Metz & Whishaw, 2002) for three trials.

#### *Ex vivo* two-photon calcium imaging

##### Brain slice preparation

Brain slices preserving the DMS and the DLS with their cortical inputs coming from somatosensory and cingulate cortex respectively were prepared as previously described. Animals were anesthetized with isoflurane before extraction of the brains. Brain slices (300 μm) were prepared using a vibrating blade microtome (VT1200S, Leica Microsystems, Nussloch, Germany). Brains are sliced in a 95 % CO_2_ and 5 % O_2_-bubbled, ice-cold cutting solution containing (in mM) 125 NaCl, 2.5 KCl, 25 glucose, 25 NaHCO_3_, 1.25 NaH_2_PO_4_, 2 CaCl_2_, 1 MgCl_2_, 1 pyruvic acid, and then transferred into the same solution at 34°C for one hour and then moved to room temperature.

##### Two-photon calcium imaging

Genetically-encoded Ca^2+^ indicator GCaMP6f was used to measure the Ca2+ activity of the neuronal somas in the striatum. GCaMP6f was expressed with recombinant AAVs injected in the DMS or the DLS. Two-photon calcium imaging was performed at λ= 940 nm with a TRiMScope II system (LaVision BioTec, Germany) using a resonant scanner, equipped with a 20x/1.0 water-immersion objective (Zeiss) and coupled to a Ti:Sapphire laser (Chameleon Vision II, Coherent, >3W, 140 fs pulses, 80MHz repetition rate). The average power of the laser emitted was set at ∼40-50mW on sample. Fluorescence is detected with a GaAsP detector (Hamamatsu H 7422-40). Scanning and image acquisitions were controlled with Imspector software (LaVision BioTec, Germany) (15.3 frames per second for 1024 x 1024 pixels, between 50 to 150 µm underneath the brain slice surface, with no digital zoom). Typical images window for calcium imaging of wide field is 392 µm x 392 µm. Cortically-evoked activity was recorded in response to electrical stimulations applied with a bipolar electrode (MicroProbes, USA) placed in the layer 5 of the cingulate cortex as previously described (Badreddine et al., 2022; Fino et al., 2018).

### DATA ANALYSIS

#### Behavior

##### Accelerating rotarod

The time to fall (latency) from the accelerating rotarod was recorded to measure the performance of the animals. The latency to fall curves (50 trials) were fitted by a one-phase association model (see (Li & Spitzer, 2020) with the equation Y=Y0 + (Ylim – Y0)*(1-exp(-K*x)) and the values of Y0, Ylim (plateau in Fig. 3) and K (slopes in Fig. 1) were compared between the tested groups. In addition, a learning index (LI) was computed by subtracting the first 2 trials from the last 2 trials, with LI = Trials_9,10_ - Trials_1,2_, in accordance with previous studies (Badreddine et al., 2022; Buitrago et al., 2004; H. H. Yin et al., 2009). For late training, learning index corresponds to a subtraction of the early trials of Day 1 from the late trials of Day 7.

##### Fixed-speed rotarod

The time to fall (latency) from the rod was measured for each tested speed. The average of the five trials per speed was calculated for each mouse.

##### Open-field

Total distance travelled during the 60 min recordings was averaged per animal. A central zone of 25×25 cm was drawn and the time spent in the center was averaged per animal. The average speed was calculated as the distance over time. The data were expressed as the average over the 60 min recording.

##### Horizontal ladder

Motor coordination was measured by using the foot fault scoring system (Metz & Whishaw, 2002) and measuring the latency to complete the task. The scoring system relies on the type of paw placement on the rungs (total miss, deep slip, slight miss, replacement, correction, partial placement). The coefficient for each of these paw placement errors is detailed in the study by Metz & Whishaw, 2002. The data were expressed as the average of the three trials of the third day.

#### Calcium imaging analysis

GCaMP6f fluorescence signals were analysed with custom-built procedures using R3.5.2 in RStudio environment as previously described and with available codes (Badreddine et al., 2022). Briefly, from manually selected ROIs in FIJI software, mean grey values and (x, y) coordinates were extracted for each ROI/slice. Calcium recordings were 700-1000 frames long and included ∼7 stimulations of cortical afferents. x(t) was the averaged intensity values of pixels in the ROI at time t for one cell. ΔF/F is obtained using y(t) = (x(t) - x0) / x0, where x0 is the mean value of the 50 % lowest values in the last 10 s. ΔF/F was then filtered with a Savitsky-Golay filter of order 3 on sliding windows of 7 frames (0.458 s). For each cell, the amplitude of response to cortical stimulations was calculated by averaging ΔF/F for 5 responses (stimulations #2 to #6). To this aim, fluorescence signals were extracted for each cell on windows of 40 frames (2.6 s) centered on the time of the first maximal amplitudes of ΔF/F (peaks) detected on cells after stimulus. Within this time window, for each response, first the peak of response was detected and then going back to the slope abrupt change between baseline and beginning of the rise, the start was detected. The amplitude of response was measured as a Delta between the values from the start and the peak points. A cell was defined as active if its amplitude of response was above a threshold defined as M+2SD, with mean (M) and standard deviation (SD) calculated individually for each neuron through the whole recordings; below this threshold, the cell was considered as inactive. The color-coded functional maps were extracted with the measure of each cell amplitude within the field of view and inactive cells are represented in white. We distinguished responses from SPNs and other cell types of striatal neurons thanks to a cell-sorting method based on calcium responses we previously developed (Becq et al., 2019).

To extract the highly active (HA) cells population, we used two different methods: (1) a thresholding using the average of the mean amplitude in naive animals to normalize the activity throughout the training conditions, i.e. for each frequency of stimulations (5 and 20 Hz) and for each territory (DMS and DLS); (2) a k-means analysis to perform amplitude clustering based on the x,y position of each cell within the field, each animal being considered independently (See Badreddine et al., 2022 for details). The group containing the cells with the maximal amplitude was defined as the HA cells and contains M ± SEM % of cells in the slice (field of the slice). The cluster area was computed using the convex hull formed by the HA cells forming the cluster determined with k-means analysis. The cluster area percentage was obtained as the HA area relative to the total area of the field.

#### Statistical analysis

The data are presented and plotted as values ± SEM, with SEM standard error of the mean, unless otherwise stated. p-values are represented by symbols using the following code: * for p< 0.05, ** for p< 0.01, *** for p< 0.001. Exact p-values and statistical tests are stated in the figure legends or in the core of the manuscript. Statistical analysis is performed using Prism 7.0 et 8.0 (GraphPad, San Diego, USA) or R environment. The sample size for the different sets of data is mentioned in the respective figure legends. Normality of each data set is checked using D’Agostino and Pearson’s test. Statistical significance was assessed using Student’s t-test or Mann-Whitney’s U-test and Wilcoxon’s signed rank test for unpaired and paired data, respectively. Pearson correlation was used for relationship between cluster size and learning index. Two-way Anova followed by Bonferroni *post-hoc* test was used to compare learning curves in WT and Hdh^CAG140/+^ groups.

## Author contribution

NB, SA, FS and EF provided the conceptual framework for the study. EF conceived the project and supervised the study; EF, SA, NB and FA designed experiments; NB, FA and EF performed experiments; NB, FA, GB and EF performed analysis; FS, SA and EF acquired funding; NB and EF wrote the manuscript; all authors read and edited the manuscript.

## Acknowledgments

The authors would like to thank the Photonic Imaging Center, PIC-GIN facility and the INMED and GIN animal facilities for mouse care. The authors would like to thank N. Tremblay, L. Galvan for helpful discussions, careful reading of the manuscript and constructive feedback. This work was supported by grants from Neuroglia (EF), Agence Nationale de la Recherche: ANR-15-IDEX-02 NeuroCoG in the framework of the *Investissements d’Avenir* program (PhD fellowship for N.B.; EF and SA), ANR-19-CE37-0026-01 ProMeSS (EF) and ANR-18-CE16-0009-

01 AXYON (FS), Fondation pour la Recherche Médicale (FRM, DEI20151234418, FS).

